# Screening and characterization of 133 physiologically-relevant environmental chemicals for reproductive toxicity

**DOI:** 10.1101/2024.03.22.584808

**Authors:** Gurugowtham Ulaganathan, Hui Jiang, Noah Canio, Ashwini Oke, Sujit Silas Armstrong, Dimitri Abrahamsson, Julia R. Varshavsky, Juleen Lam, Courtney Cooper, Joshua F. Robinson, Jennifer C. Fung, Tracey J. Woodruff, Patrick Allard

## Abstract

Reproduction is a functional outcome that relies on complex cellular, tissue, and organ interactions that span the developmental period to adulthood. Thus, the assessment of its disruption by environmental chemicals is remarkably painstaking in conventional toxicological animal models and does not scale up to the number of chemicals present in our environment and requiring testing.

We adapted a previously described low-throughput *in vivo* chromosome segregation assay using *C. elegans* predictive of reproductive toxicity and leveraged available public data sources (ToxCast, ICE) to screen and characterize 133 physiologically-relevant chemicals in a high-throughput manner. The screening outcome was further validated in a second, independent *in vivo* assay assessing embryonic viability. In total, 13 chemicals were classified as reproductive toxicants with the two most active chemicals belonging to the large family of Quaternary Ammonium Compounds (QACs) commonly used as disinfectants but with limited available reproductive toxicity data. We compared the results from the *C. elegans* assay with ToxCast *in vitro* data compiled from 700+ cell response assays and 300+ signaling pathways-based assays. We did not observe a difference in the bioactivity or in average potency (AC50) between the top and bottom chemicals. However, the intended target categories were significantly different between the classified chemicals with, in particular, an over-representation of steroid hormone targets for the high Z-score chemicals.

Taken together, these results point to the value of *in vivo* models that scale to high-throughput level for reproductive toxicity assessment and to the need to prioritize the assessment of QACs impacts on reproduction.

## INTRODUCTION

The identification of environmental chemicals that impact reproduction has long been challenging ^1^. The origin of this difficulty is plural. Reproduction is a remarkably complex process involving the interaction of multiple organ systems over long periods of time. The initiation of germ cell development happens early during mammalian development but only completes decades later in humans ^2^. The Organisation for Economic Co-operation and Development (OECD) validated tests, such as the OECD 416 two-generation reproduction toxicity test ^3^, and OECD 414 prenatal developmental toxicity test ^4^ can capture various windows of exposure important for reproduction but are not scalable for the screening of the large number of environmental chemicals and their mixtures registered for use worldwide, recently estimated *ca.* 350,000 chemicals ^5^. These challenges stand in stark contrast to the dire need to address the root causes of infertility. In its 2023 report, the World Health Organization reported that 1 in 6 people worldwide, or approximately 17.5% of the adult population, suffers from infertility with a lifetime prevalence of 17.8% in high-income countries and 16.5% in low- and middle-income countries ^6^.

The nematode *Caenorhabditis elegans* (*C. elegans*) is a valuable model for the rapid assessment of environmental reproductive effects. The advantages of using *C. elegans* for such studies stem from the unique combination of features exhibited by the nematode. *C. elegans* is a highly tractable model system with a large degree of conservation of the fundamental aspects of reproduction with higher organisms, including the pathways underlying the process of meiotic division which results in the generation of haploid gametes ^7,8^. Its transparency and spatially distinct meiotic stages allow for the direct visualization and easy identification of meiotic stages and meiotic defects. We previously showed that this model can be leveraged for the dissection of reproductive toxicity mechanisms as demonstrated in the context of a variety of environmental exposures such as endocrine disruptors ^9,10^, metals ^11^, pesticides ^12^, or alcohol ^13^. We also showed that genetic tools developed to identify new genes and pathways implicated in germ cell differentiation can be applied for the rapid identification of mammalian reproductive toxicants ^12,14,15^. In these studies, a transgenic reporter line which reports errors in chromosome segregation was used and validated. This line takes advantage of the naturally hermaphroditic (XX) state of *C. elegans* and the naturally low occurrence of males (<0.2%). The appearance of males in the population is often a result of meiotic segregation errors, *i.e.* errors during the segregation of the X chromosome ^16,17^. Therefore, the screen utilizes a reporter strain (*Pxol1::GFP*) in which GPF expression is under the control of a male-specific promoter that is expressed early during embryogenesis allowing the direct visualization of errors in segregation of the X chromosome which generates male embryos. This screen, termed the Green Eggs and HIM (*high incidence of males)* screen ^16^, when applied to environmental chemicals, showed notable predictivity towards mammalian reproductive endpoints, with a 69% maximum balanced accuracy ^14^.

Here, we applied this assay to the reproductive toxicity assessment of a chemical database of 133 chemicals compiled from a suspect screening, non-targeted analysis of blood samples from pregnant individuals in San Francisco, combined with other chemicals of high concern for human health impact such as alternative flame retardants (AFRs), pesticides and biocides, perfluoroalkyl and polyfluoroalkyl substances (PFAS), plasticizers, and quaternary ammonium compounds (QACs) ^18,19^. The screen identified 13 chemicals with suspected reproductive toxicity. Amongst the 5 most potent chemicals that were further tested for their impact on reproduction, 4 showed a significant increase in embryonic lethality. The mining of US EPA’s Toxicity Forecaster (ToxCast) and of the National Toxicology Program Interagency Center for the Evaluation of Alternative Toxicological Methods’ Integrated Chemical Environment (ICE) databases raised significant concerns about *in vitro* and *in vivo* data paucity and relevance for reproductive toxicity of chemicals that are commonly used in consumer products.

## MATERIAL AND METHODS

### C. elegans strains

*C. elegans* strains were cultured as previously described at 20°C on nematode growth medium (NGM) plates ^20^. The N2 Bristol strain was used as the wild-type strain and the following transgene was used in this study: LGV, yIs34[Pxol-1::GFP, rol-6]. All chemicals used in the study were acquired from Sigma Aldrich.

### Blinding, exposure, growth conditions, and image acquisition

All chemicals obtained by our laboratory were independently coded by the UCSF team and provided to the UCLA team encoded. Thus, all following experiments were performed without knowledge of chemical identity. Chemical information was revealed following completion of the screening phase of the project.

For exposure in deep-well 96-well plates, we first generated a large population of L1 larvae. To this effect, gravid worms from NGM plates were collected by washing the plates with 1-2 mL of M9 and then transferring the worms to a 15 mL conical tube. The gravid worms were left to sediment to the bottom of the 15 mL conical tube for approximately 5-10 min. The supernatant was removed without disturbing the worm pellet. We added 1 mL of standard bleach/NaOH solution into the tube and incubated for 2-3 minutes at room temperature. We monitored the progress of the reaction under a dissecting microscope to confirm that all the worms had stopped moving. The worms were collected by centrifuging for 1 min at 3,000 rpm (900g). We removed the supernatant and added 4 mL of sterile M9 to neutralize the reaction. The worms were washed twice by centrifuging for 1 min at 3,000 rpm (900g). Then the supernatant was removed and replaced with M9 up to 200 µL. Using a glass Pasteur pipette, a drop of the worm/M9 mix was added to the center of clear NGM plates that were incubated for ∼24 hours at 20 °C to allow the embryos to hatch. After 24 hours, hatched L1 larvae were collected by washing them with 1-2 mL of M9. After assessment of worm concentration under a stereomicroscope, 60,000 L1 larvae were added to a 600ml culture flask with 6 ml of a stock OP50 solution (100mg/ml). M9 was added to adjust the total amount of solution in the 600ml flask to 60ml. The worms were then incubated at 20 °C on a shaker at 150-200 rpm until the worms reached the L4 stage.

The L4 larvae were transferred from a 600 ml flask to 2 conical 50 mL tubes filled with M9 buffer and the L4s were left to settle for approximately 5-10 min. This step was repeated 2 times in order to clear the L4s from potential embryos or debris. After the last wash, the worms were resuspended in 20 ml and their concentration was adjusted to 100 L4s per 50 µl. The worms were then transferred to a 96 deep-well plate (Corning Axygen™ PDW20CS) with 50 µl of L4 solution (i.e. 100 L4s) and 350 µl of OP50 solution (5.7mg/ml). Then, 0.5 µL of either the test chemical dissolved in DMSO, which was also used as a negative control, or Nocodazole (positive control) were added to the desired wells. Experimental repeats of the chemicals were performed in two distinct plate layouts where well position was changed to minimize the impact of edge effect. For all wells, the final DMSO concentration was 0.1%. The plates were sealed using adhesive films, wrapped with aluminum foil and transferred to a shaker (180 rpm) for 24 hours.

A total of 133 chemicals were tested. First, 111 chemicals were tested at two concentrations (30 μM and 100 μM) based on the good predictive value of the screens at the highest concentration ^21^, with four biological repeats each performed in experimental duplicates. Of these 111 chemicals, a subset of 61 chemicals were further tested at all four concentrations (10 μM, 30 μM, 50 μM and 100 μM) with five biological repeats performed in duplicate in order to facilitate future comparison with a comparable yeast assay ^22^ (**Table S1**). Due to solubility limitations, 21 chemicals were tested at lower concentrations than those indicated (**Table S1**). One chemical, Butylparaben (CAS 94-26-8) was tested but not included in the results since its exposure led to high levels of background fluorescence in the worms.

### Image acquisition

After washing, the worms were transferred to the wells of a black wall/clear bottom 384-well plate (Greiner Bio-One 781986). Using a multichannel pipette, 10 µL of levamisole (50 µM) was added to each well followed by a 20-minute incubation. For image acquisition, the plates were transferred to an ImageXpress imaging platform and microscopic data was captured using the Meta Xpress software. Over 6,000 images were analyzed for the outcome of interest, i.e. GFP+ embryos. Wells with high proportions of dead worms (> 10%) or with nematodes showing ectopic reporter expression, such as in the spermatheca or in the pharynx, were excluded from the counts. Due to the complexity of identifying positive embryonic events within the context of natural background autofluorescence emitted by the intestinal tissue ^23,24^ as well as strong ectopic fluorescence noticed in the pharyngeal tissue for some chemicals, we employed a manual review of each captured image, with two independent scorers of the image datasets.

### Chemical ranking methodology

We sorted all the observed GFP+ counts for the chemicals into their four concentration exposure groups: 10 μM, 30 μM, 50 μM, and 100 μM. Within each concentration group, we calculated the ratio of worms containing at least one GFP-positive event to the total number of worms for each replicate. Next, we averaged the biological repeats for a given chemical within a tested concentration. After obtaining the ratio values, we calculated the Z-score for each chemical using the following formula: z = (x-μ)/σ, where x is the averaged ratio of the biological repeats as described above, μ is the mean for all chemicals within each concentration, and σ is the corresponding standard deviation. For our ranking methodology, chemicals with a Z>1 1 (*i.e.* chemicals greater than 1 standard deviation away from the mean) at any of the four tested concentrations were categorized as high Z-score chemicals. Conversely, chemicals that consistently exhibited Z-scores <1 across all tested concentrations were labeled as low Z-score chemicals.

### Embryonic lethality assessment

The top 5 chemicals from the high Z-score category and the bottom 5 chemicals from the low Z-score category were tested for their ability to elicit embryonic lethality. Embryonic lethality was performed three times for each exposure. L4 worms were exposed to the selected chemicals at 100 μM for 24 hours. Following exposure, a total of 10 worms per condition were individually transferred to one small NGM plate (35 x 10mm) each, in triplicate. The worms were left to lay approximately 100 eggs on each plate after which the worms were removed, and the exact number of embryos recorded. After 2 days at 20°C, the number of larvae that hatched from these embryos was recorded for each plate.

### ToxCast dashboard data mining

Following our chemical group categorization, we compared our data with the EPA’s Toxicity Forecaster (ToxCast) dashboard. This dashboard houses a comprehensive collection of high-throughput *in vitro* assay screening and exposure data for thousands of environmental chemicals. For our study’s integrated analysis, we employed an approach that considered three measures: (1) Bioactivity: This measure utilized the ‘Hit Call’ data, where we calculated the ratio of active assay hits as listed in the ToxCast dataset to the total number of assays for a given chemical. This metric was used as a representation of the bioactivity of the chemicals identified in our *in vivo* screen. (2) Log10 AC50 values: from the *in vitro* assay data, we transformed the AC50 values into their Log10 values (Log10 AC50). For chemicals that had assays without any associated AC50 values, we assigned an arbitrary Log10 value of ‘3’ as previously described ^14^. (3) Biological/Signaling endpoints: For this measure, we used the ‘Intended Target Family’ data filtered from active hit call data in the ToxCast data. The counts and distribution of the different target families were analyzed for each chemical to identify *in vitro* endpoints that may be associated with the outcome of the worm screen. Intended target families in ToxCast included two non-descriptive categories “channel 1” and “channel 2” which represent a collection of various *in vitro* assays where fluorescence signal is captured through two different channels. These categories were included in the bioactivity (Log10 AC50) comparison but were excluded when specifically examining the differential representation of intended target families between chemical groups. One chemical from the high Z-score group, tri-p-tert-butylphenyl phosphate (CAS# 78-33-1) does not have bioactivity data available through the ToxCast dashboard and therefore was not included in these analyses.

### Integrated Chemical Environment (ICE)

We employed the ‘Chemical Characterization’ tool from the Integrated Chemical Environment ICE v4.0.1 database (https://ice.ntp.niehs.nih.gov/), which is a collection of curated data from NICEATM and other toxicological sources (Tox21 consortium), to perform Principal Component analysis comparison, and identify product use categories of the highest and lowest Z score chemicals using their CAS ID numbers.

### Statistical analyses

All statistical analyses of the ToxCast data were performed on RStudio version 2022.02.3.

## RESULTS

### Identification of reprotoxic environmental chemicals through the *in vivo C. elegans* aneuploidy screening platform

In the present study, we expanded a previously described low-throughput screening *in vivo* assay using *C. elegans* ^14^ into a high-throughput/high-content platform to simultaneously rapidly assess 100s of chemicals at multiple doses (see material and methods section). Briefly, this approach uses a combination of deep-well 96-well plates for exposure and a 384-well plate based-high-content imaging platform to assess ∼100 nematodes per well. We collected a total of 6,000 data points for 133 chemicals in 4-5 biological repeats and 2 experimental replicates (**Figure 1, Table S1**).

**Figure 1:**
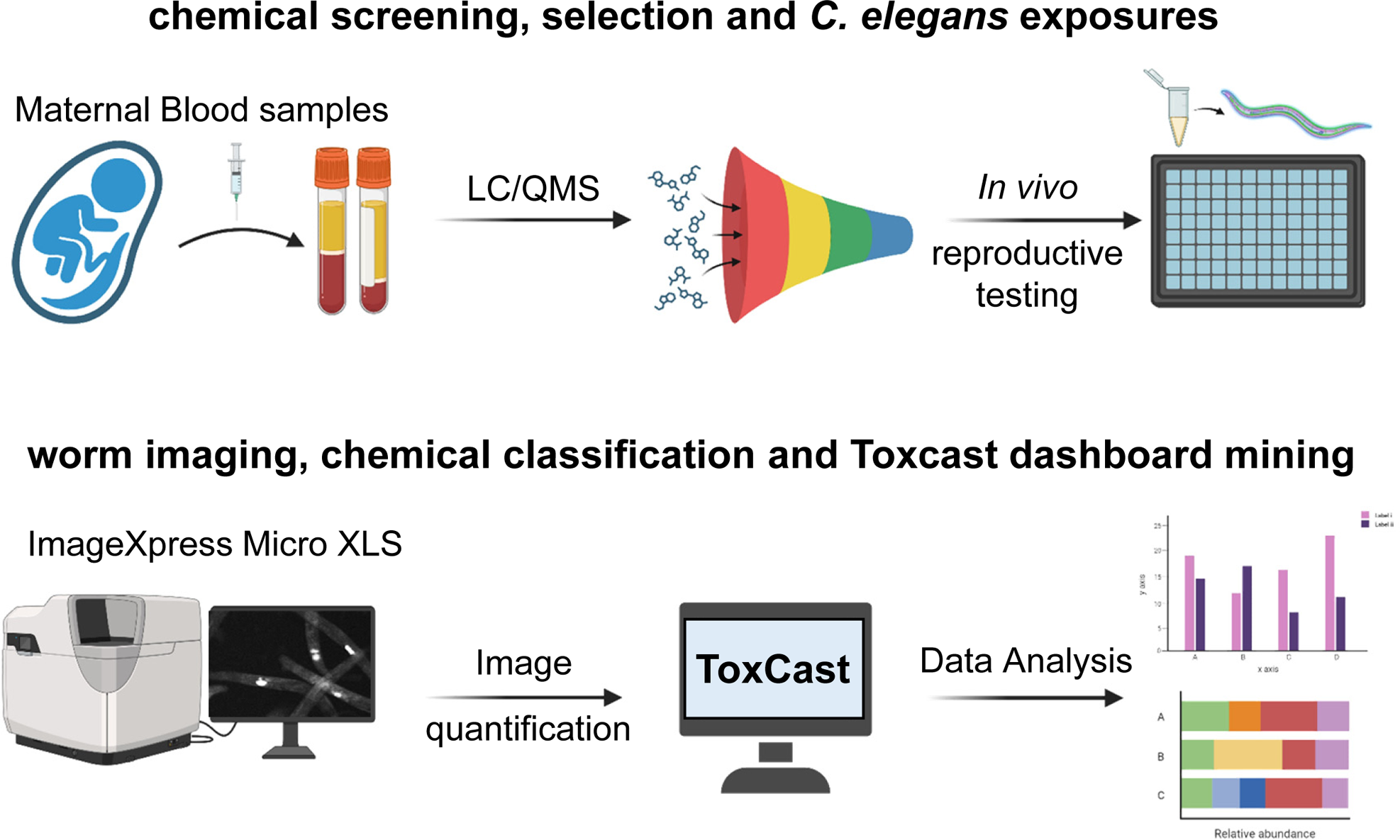
Overview of study methodology. A chemical library composed of chemicals identified in maternal serum samples supplemented with chemicals of potential reproductive significance is tested in a set of *in vivo* reproductive assays in *C. elegans*. The aneuploidy (X chromosome segregation error) reporting worm strain containing yIs34[*Pxol-1::GFP, rol-6*] was exposed in 96-well plates for 24 hours. GFP expression was visualized using the ImageXpress Micro XLS imaging platform. After image acquisition, GFP+ events were quantified, and chemicals were ranked by Z-score. A subset of these chemicals were further tested for embryonic viability. The *C. elegans* outcome was compared to the *in vitro* ToxCast assay data to correlate *in vitro* bioactivity with reproductive toxicity in *C. elegans*.

To accurately report and rank chemicals’ potency, we calculated chemicals by Z-score ^25^ within each concentration group. We used the robust metric of Z-score (see material and method section) to identify chemicals that exhibited a Z-score of at least 1 (*i.e*. > 1 standard deviation greater than the mean) at any of the four concentrations **(Table S1, Fig 2A)**. These chemicals were labeled as high Z-score chemicals, and across all tested concentrations, 13 such chemicals fit this criterion (**Table 1**). Chemicals consistently displaying the lowest or low z-scores (i.e. < 1) at all concentrations were classified as low Z-score chemicals. A representative side-by-side comparison of chemicals from these two groups at a dose of 100 μM, namely methylbenzethonium chloride (high Z-score group) and monobenzyl phthalate (low Z-score group) is shown in **Figure 2B**, revealing a noticeable difference in presence of GFP+ embryos within the body of exposed worms. These were comparable to the positive and negative controls used in our study, the microtubule disrupting agent nocodazole, and DMSO, respectively (**Figure 2B**).

**Figure 2:**
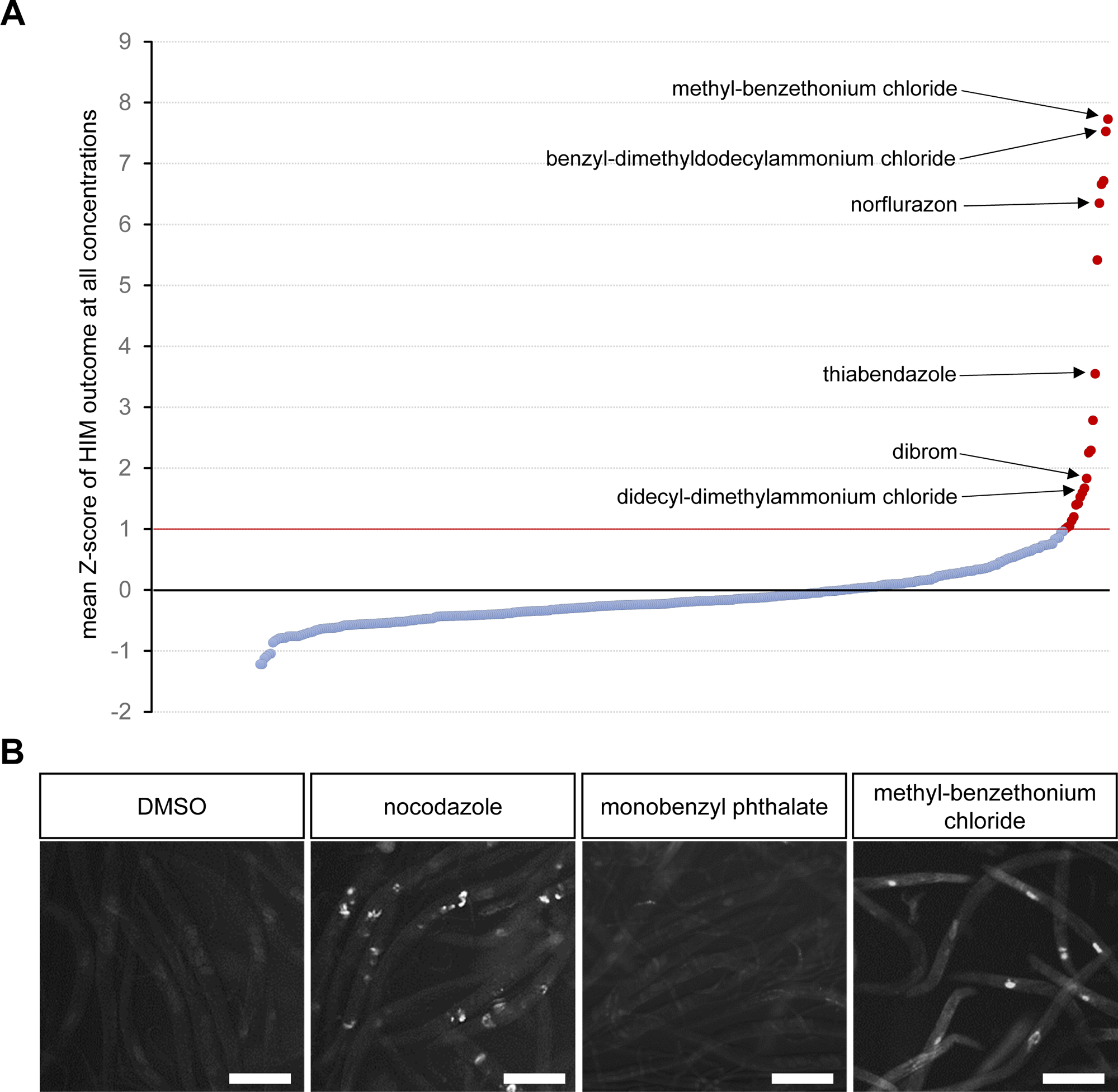
Identification of aneuploidy-inducing chemicals. (A) Visualization of chemical ranking by Z-score at all concentrations. The red bar represents the Z-score cut-off of 1 above which chemicals were classified as high Z score chemical. (B) Observation of GFP+ embryos in 0.1% DMSO (negative control), nocodazole (positive control), monobenzyl Phthalate (Z-score <1) and methylbenzethonium chloride (Z-score >1) all at 100 μM. Error bars = 200 μm.

**Table 1:**
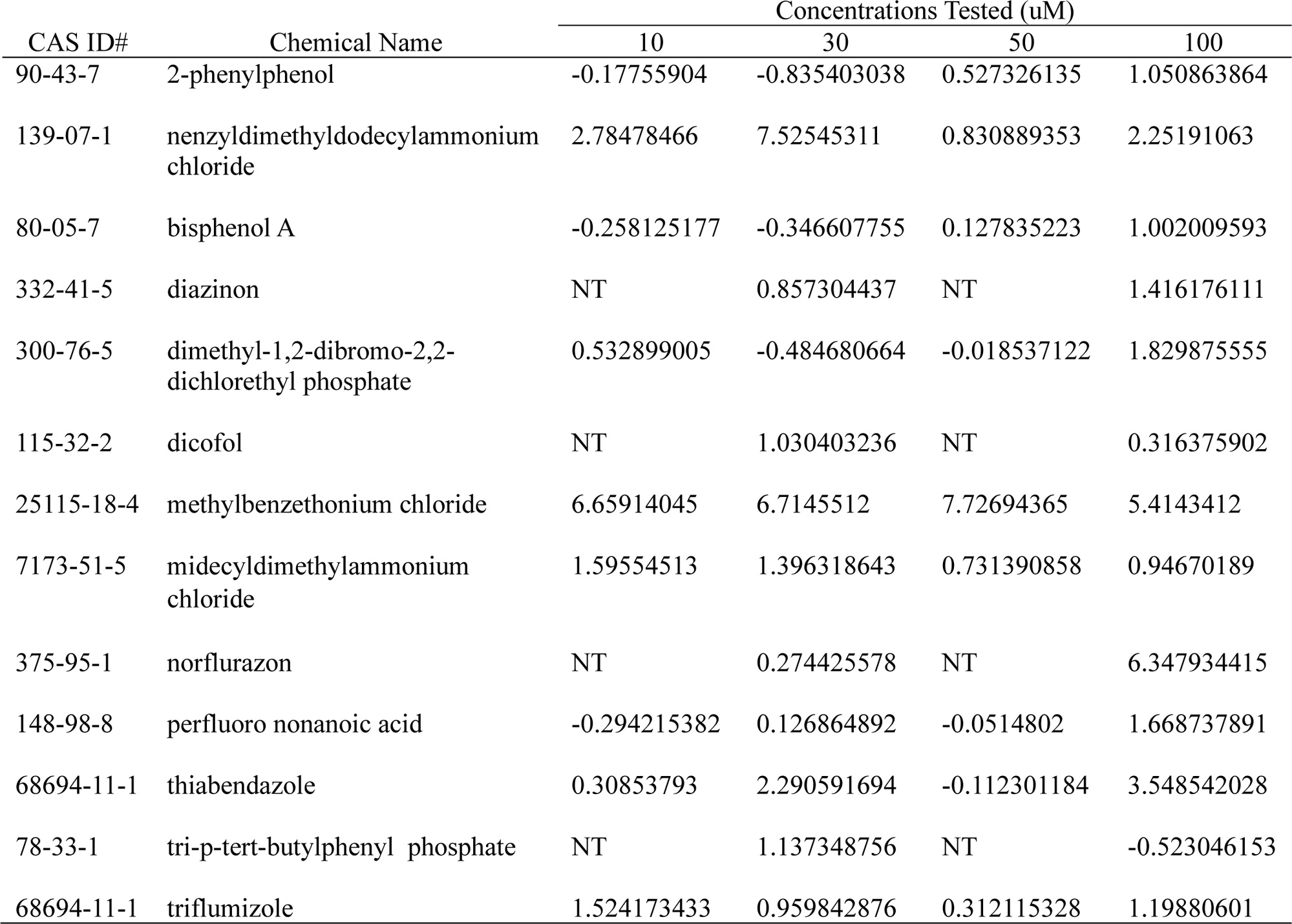
Chemicals with a Z score>1 at any tested concentration.

Genetically- or chemically-altered germline function is often associated with embryonic lethality ^9,12,14,26^. Thus, we selected the top 5 and bottom 5 ranked chemicals by Z-score (**Tables 1, 2**) and assessed their impact on embryonic lethality (see Material & Methods section). From the top 5 chemicals, 4 of them showed a statistically significant increase in embryonic lethality compared to DMSO control (Brown-Forsythe and Welch ANOVA test with Dunnett correction), with the top two chemicals (methylbenzethonium chloride and benzyldimethyldodecylammonium chloride) displaying a remarkable 20% mean embryonic lethality at 100 μM (**Figure 3A**). Conversely, none of the bottom 5 chemicals were positive for embryonic lethality (**Figure 3B**). Collectively, these findings indicate that this *in vivo* screening platform identified a specific set of chemicals carrying reproductive toxicity.

**Figure 3:**
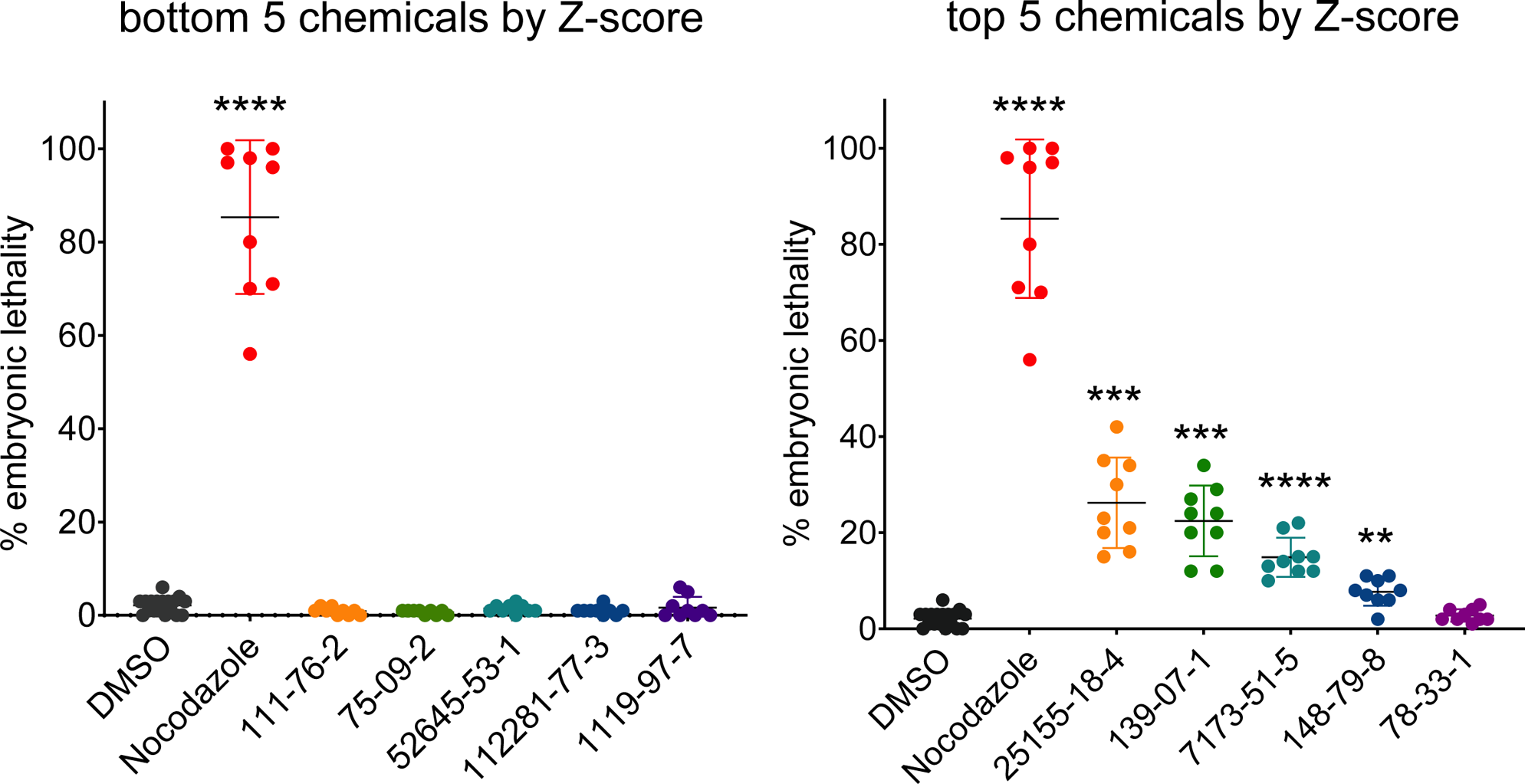
Embryonic lethality scores for the top and bottom ranked chemicals. Embryonic lethality outcome for top 5 chemicals (Z-score >1) vs. bottom 5 chemicals (Z-score <1) as identified in the *in vivo* aneuploidy screen. Chemicals were tested together at 100 μM and 0.1% DMSO and 100 μM nocodazole were used as negative and positive controls respectively. N=9, 10 worms each. Adjusted P values ****P < 0.0001, ***P < 0.001, **P < 0.01, Brown-Forsythe and Welch ANOVA test with Dunnett correction for multiple comparisons. For chemical names from CAS ID# refer to Tables 1 and 2.

**Table 2:** Chemicals with a Z score <1 at all tested concentrations.

| CAS ID# | Chemical Name | Concentrations Tested (uM) |  |  |  |
| --- | --- | --- | --- | --- | --- |
|  |  | 10 | 30 | 50 | 100 |
| 111-76-2 | ethylene glycol butyl ether (EGBE) | -0.493188189 | -0.760018538 | -0.428279589 | -0.786605971 |
| 3194-55-6 | hexabromocyclodecane (HBCD) | -0.532066923 | -0.637397676 | -0.262285387 | -0.635415477 |
| 121-75-5 | malathion | -0.57865188 | -0.633130582 | -0.195971923 | -0.080772843 |
| 75-09-2 | methylene Chloride | -0.760359221 | -0.205061778 | -0.145673888 | -0.320872732 |
| 2528-16-7 | monobenzyl phthalate | -0.384659645 | -0.081544854 | -0.425217575 | -0.614814203 |
| 335-76-2 | perfluorodecanoic acid | -0.760359221 | -0.402192131 | NDA | -1.219646999 |
| 52645-53-1 | permethrin | -0.55800923 | -0.735099931 | -0.198504842 | -0.561941711 |
| 175013-18-0 | pyraclostrobin | -0.436371926 | -0.067060827 | -0.242144092 | -0.523376499 |
| 302-79-4 | retinoic Acid (RA) | -0.654868704 | -0.096655283 | -0.429698571 | -0.586138019 |
| 13674-87-8 | tris(1,3-dichloro-2-propyl) phosphate (TDCIPP) | -0.56212385 | -0.236069063 | -0.422525695 | -0.350506139 |
| 112281-77-3 | tetraconazole | -0.369829188 | -0.574755817 | -0.344775632 | -1.124919234 |
| 1119-97-7 | tetradonium bromide | -0.760359221 | -0.166809423 | -0.486457849 | -0.864417879 |

#### Integration of high-throughput in vitro ToxCast data

To integrate the output of our screen with publicly accessible high-throughput *in vitro* toxicity data, we mined the Toxicity Forecaster (ToxCast) dashboard. We particularly aimed to compare our pre-classified datasets (top and bottom-ranked chemicals by Z-score) with multiple variables of interest: number of positive assays (*i.e.* hit calls), AC50 values, and intended targets associated with the hit calls.

We first assessed all tested assays for a given chemical. For these assays, we calculated the bioactivity ratio, a value derived from the ratio of the active hit calls (positive assays) by the total number of assays in which the chemical was tested in the ToxCast dataset (**Tables 3, 4**). This value can therefore serve as a general measure of chemical bioactivity. Interestingly, for high Z score and low Z score chemicals in the *C. elegans* assay, no obvious difference in bioactivity was observed (**Figure 4**). Hence, a comparison of the average bioactivity values for the high Z score and low Z score groups did not reveal a significant difference between the two groups (high Z score group = 0.249 and low Z score group = 0.234, P=0.46 by unpaired t test with Welch’s correction). For instance, within the high Z score chemicals for the worm assay, a chemical such as methylbenzethonium chloride displayed a bioactivity ratio of 0.489, in contrast to the more modest value of 0.086 associated with another high Z score chemical: thiabendazole. Similarly, for the low Z score chemicals in the worm assay, tetraconazole exhibited a ratio of 0.333 while monobenzyl phthalate had a ratio of 0.026. We extended our analysis to include the AC50 values of the chemical’s assays to further substantiate this finding. The visualization of Log10 converted AC50 values corroborated the findings from the bioactivity ratio: the Log10 AC50 values were equally spread across the X axis for both chemical groups without discernible pattern (**Figure 5**).

**Figure 4:**
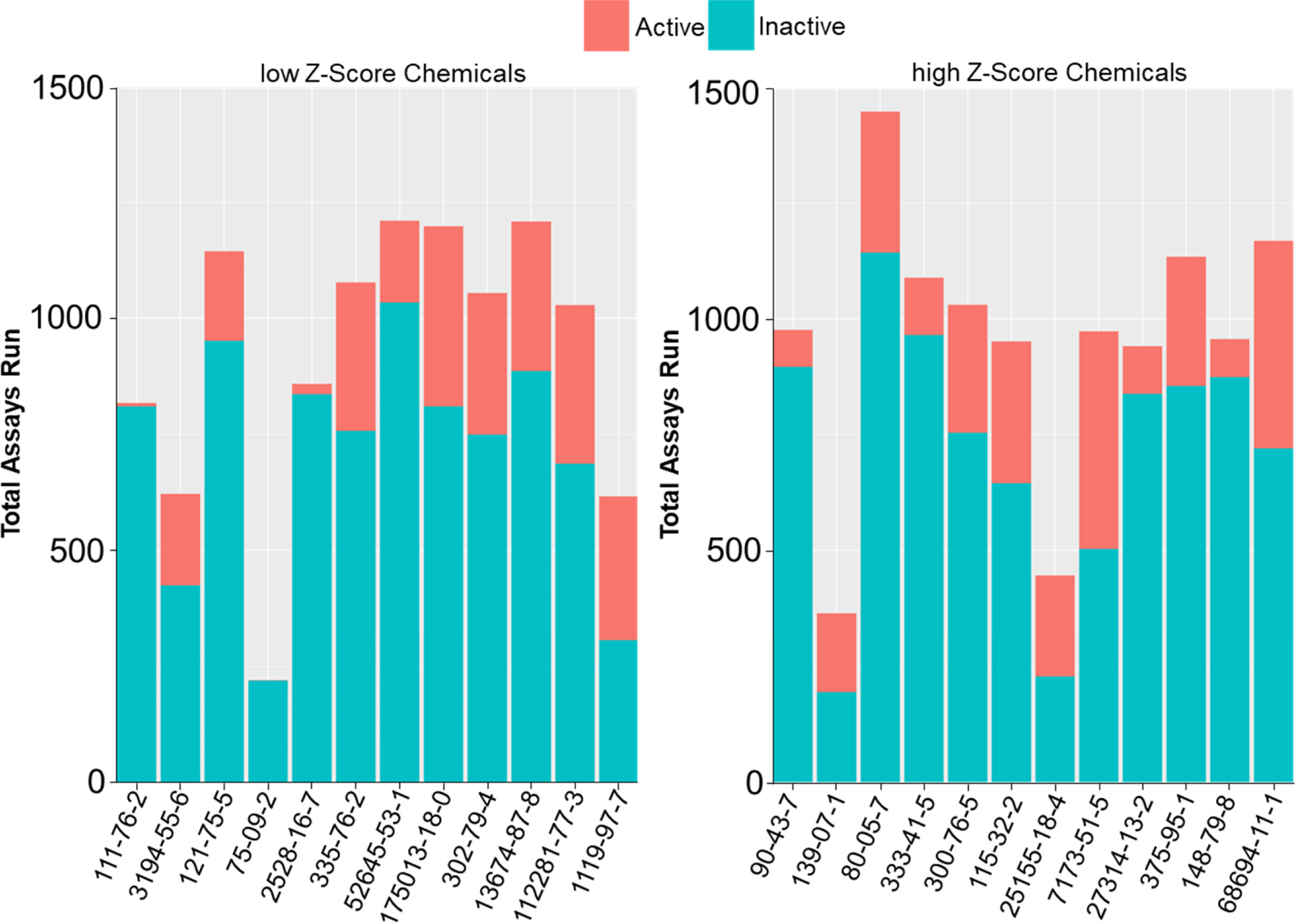
Active and inactive assays from the *in vitro* ToxCast data. Total number of assays from the ToxCast dashboard shown as a sum of the active and inactive assay hits for the top and bottom chemicals classified by Z-score. For chemical names from CAS ID# refer to Tables 1 and 2.

**Figure 5:**
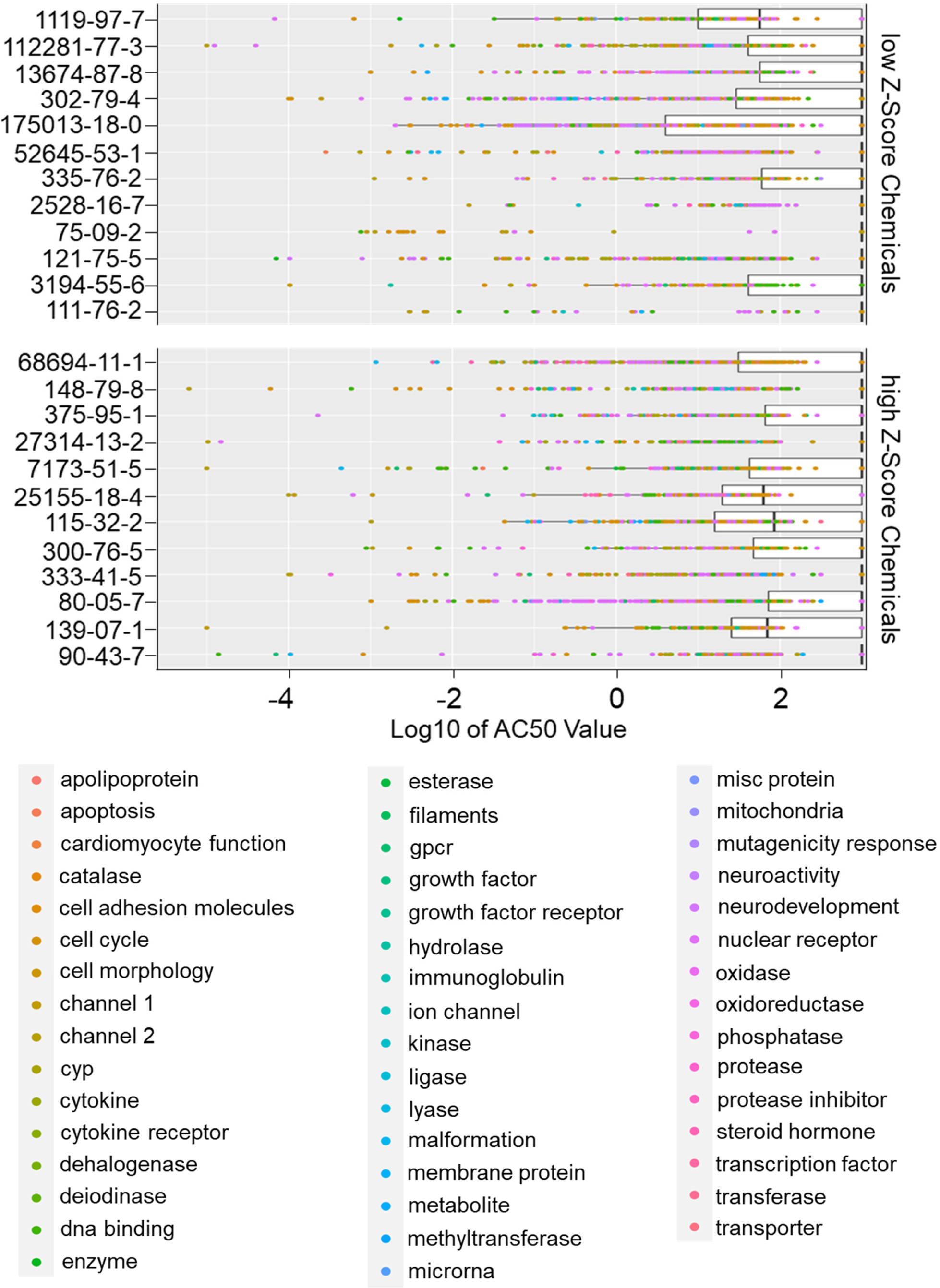
Bioactivity of the top and bottom chemicals. Distribution of Log10 AC50 values between bottom and top chemicals from the *C. elegans* aneuploidy assay. The box-plots are ordered by chemical ranking from Low Z-score chemicals to high Z-score chemicals.

**Table 3.** Bioactivity ratios of high Z-score chemicals.

| Chemical Name | Active hit calls | Total assays | Bioactivity ratio |
| --- | --- | --- | --- |
| 2-Phenylphenol | 80 | 976 | 0.082 |
| Benzyldimethyldodecylammonium chloride | 170 | 364 | 0.467 |
| Bisphenol A | 304 | 1447 | 0.210 |
| Diazinon | 124 | 1089 | 0.114 |
| Dimethyl-1,2-dibromo-2,2-dichlorethyl phosphate | 276 | 1030 | 0.268 |
| Dicofol | 306 | 951 | 0.322 |
| Methylbenzethonium chloride | 218 | 446 | 0.489 |
| Didecyldimethylammonium chloride | 470 | 973 | 0.483 |
| Norflurazon | 103 | 941 | 0.109 |
| Perfluoro nonanoic acid | 279 | 1134 | 0.246 |
| Thiabendazole | 82 | 956 | 0.086 |
| Triflumizole | 448 | 1168 | 0.384 |

Note: tri-p-tert-butylphenyl phosphate (CAS# 78-33-1) does not have bioactivity data available through the ToxCast dashboard and is therefore not listed.

**Table 4.** Bioactivity ratios of low Z-score chemicals.

| Chemical Name | Active hit calls | Total assays | Bioactivity ratio |
| --- | --- | --- | --- |
| Ethylene glycol butyl ether (EGBE) | 7 | 817 | 0.009 |
| Hexabromocyclodecane (HBCD) | 197 | 621 | 0.317 |
| Malathion | 193 | 1144 | 0.169 |
| Methylene Chloride | 1 | 220 | 0.005 |
| Monobenzyl phthalate | 22 | 858 | 0.026 |
| Perfluorodecanoic acid | 320 | 1077 | 0.297 |
| Permethrin | 176 | 1210 | 0.145 |
| Pyraclostrobin | 388 | 1198 | 0.324 |
| Retinoic Acid (RA) | 305 | 1054 | 0.289 |
| Tris(1,3-dichloro-2-propyl)phosphate (TDCIPP) | 322 | 1208 | 0.267 |
| Tetraconazole | 342 | 1028 | 0.333 |
| Tetradonium bromide | 310 | 616 | 0.503 |

We then focused our analysis on the classification by intended molecular target of all assays with active hit calls. We identified 43 distinct target families with active hits in response to either chemical group. Amongst these, several families were more prominently represented than others: cell cycle (N = 1106), nuclear receptor (N = 849), DNA binding (N = 482), cytokine (N = 418), neurodevelopment (N = 351) and cell adhesion (N = 194). In terms of specific chemicals, didecyldimethylammonium chloride had the highest number of active hits targeting families like cell cycle (N = 113), followed by thiabendazole (N = 100) in the positive group **(Figure 6).**

**Figure 6:**
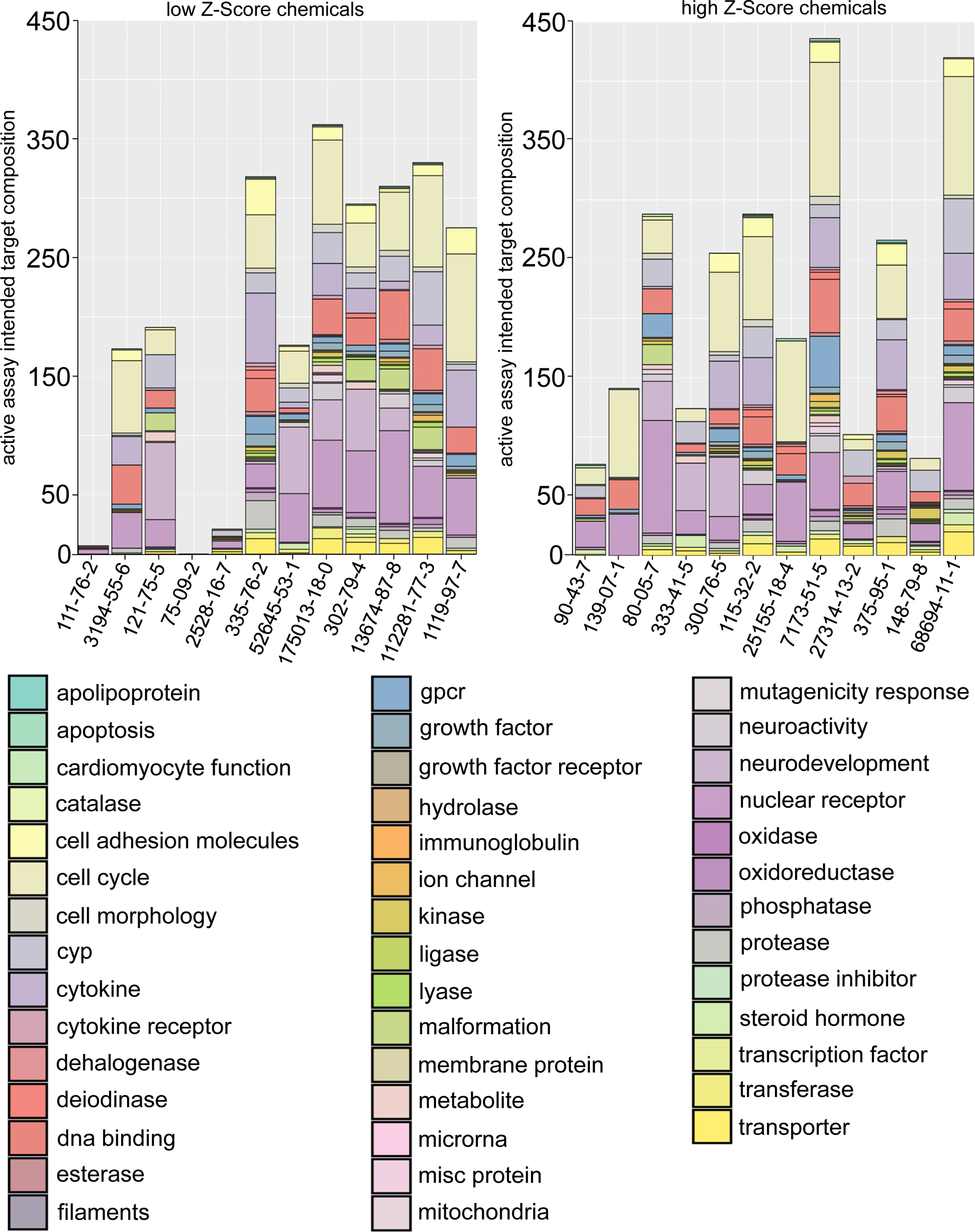
Classification of intended target families. All assays with active hit calls from the bottom and top chemicals in the *C. elegans* assay are visualized as a normalized percent-bar graph. Methylene chloride was excluded from this analysis since it was only positive for the not descriptive “*channel 2*” category.

We performed a categorical analysis between the high and low Z-score chemicals to establish whether a statistically significant difference existed in their intended target family responses. We also sought to identify the exact target families responsible for driving this difference. The categorical comparison of the two groups was highly significant (χ²= 1.37E-38) (**Table 5**). To identify the intended target families that may drive the difference between the top and bottom chemicals, we performed a Fisher Exact Test both with and without adjustment for multiple comparisons (**Table 5**). Without adjustment, 5 target families were significantly over-represented in the top chemicals (steroid hormone, cell cycle, GPCR, kinase, and deiodinase) while only “steroid hormone” remained significant after adjustment.

**Table 5:** Molecular target categories comparison.

| **Name** | **Active Assays**<br>**Bottom Chemical** | **Active Assays**<br>**Top Chemicals** | **Fisher Exact** | **FDR adj**<br>**FisherExact** |
| --- | --- | --- | --- | --- |
| **Steroid hormone** | 11 | 37 | 0.000431 | 0.018115 |
| **Cell cycle** | 480 | 626 | 0.007459 | 0.156631 |
| **Gpcr** | 63 | 102 | 0.011238 | 0.157328 |
| **Kinase** | 16 | 34 | 0.019903 | 0.186766 |
| **Deiodinase** | 11 | 26 | 0.022234 | 0.186766 |
| **Apoptosis** | 1 | 6 | 0.081175 | 0.568227 |
| **Dehalogenase** | 3 | 9 | 0.10123 | 0.607382 |
| **Cardiomyocyte function** | 1 | 5 | 0.135322 | 0.710441 |
| **Mitochondria** | 10 | 16 | 0.236846 | 0.945855 |
| **Esterase** | 10 | 16 | 0.236846 | 0.945855 |
| **Hydrolase** | 2 | 5 | 0.270689 | 0.945855 |
| **Neuroactivity** | 35 | 45 | 0.293081 | 0.945855 |
| **Transporter** | 67 | 81 | 0.329758 | 0.945855 |
| **Oxidoreductase** | 20 | 26 | 0.349409 | 0.945855 |
| **growth factor receptor** | 5 | 8 | 0.357916 | 0.945855 |
| **Cytokine receptor** | 10 | 14 | 0.360326 | 0.945855 |
| **Cyp** | 174 | 198 | 0.420697 | 0.979568 |
| **Nuclear receptor** | 402 | 447 | 0.492056 | 0.979568 |
| **Protease inhibitor** | 9 | 11 | 0.503453 | 0.979568 |
| **Ligase** | 4 | 5 | 0.562571 | 0.979568 |
| **Filaments** | 5 | 6 | 0.568756 | 0.979568 |
| **Mutagenicity response** | 8 | 9 | 0.584528 | 0.979568 |
| **DNA binding** | 233 | 249 | 0.663085 | 0.979568 |
| **Misc protein** | 5 | 5 | 0.683913 | 0.979568 |
| **Oxidase** | 5 | 5 | 0.683913 | 0.979568 |
| **Transcription factor** | 3 | 3 | 0.70272 | 0.979568 |
| **Transferase** | 32 | 31 | 0.743804 | 0.979568 |
| **Ion channel** | 13 | 12 | 0.74387 | 0.979568 |
| **Growth factor** | 33 | 32 | 0.744556 | 0.979568 |
| **Catalase** | 5 | 4 | 0.793523 | 0.979568 |
| **Lyase** | 16 | 14 | 0.795718 | 0.979568 |
| **MicroRNA** | 4 | 3 | 0.813002 | 0.979568 |
| **Apolipoprotein** | 4 | 3 | 0.813002 | 0.979568 |
| **Metabolite** | 21 | 18 | 0.830782 | 0.979568 |
| **Phosphatase** | 17 | 14 | 0.841617 | 0.979568 |
| **Cell morphology** | 33 | 28 | 0.877353 | 0.979568 |
| **Cytokine** | 210 | 207 | 0.88397 | 0.979568 |
| **Membrane protein** | 5 | 3 | 0.886276 | 0.979568 |
| **Protease** | 65 | 57 | 0.913858 | 0.984155 |
| **Cell adhesion molecules** | 105 | 89 | 0.971261 | 1 |
| **Malformation** | 74 | 28 | 1 | 1 |
| **Neurodevelopment** | 226 | 125 | 1 | 1 |

In sum, while a difference in the bioactivity of the two classified datasets based on our screen was overall not observed, categorical distinctions in the biological responses to these chemicals were evident *via* their differential elicitation of some intended target families.

### Mining of the Integrated Chemical Environment database

The Integrated Chemical Environment database is a collection of curated data from several diverse toxicological sources, primarily NICEATM and the Tox21 consortium ^27^. Access to the chemical characterization tool in the database allowed us to investigate potential differences in physico-chemical properties between the high and low Z-score chemicals in the *C. elegans* assay as well as the potential product categories where chemicals identified in our screen are commonly present which may therefore represent their sources of exposure. As shown in **Figure 7A**, by PCA plot of the chemicals by chemical properties (e.g. molecular weight, boiling point, KOA, etc), no clear separation was apparent. With regards to product use, compounds such as didecyldimethylammonium chloride and 2-phenylphenol that are part of the high Z-score chemical group were identified in household supplies like disinfectants and surface cleaners belonging to the cleaning products category. Likewise, within the low Z-score group, substances like ethylene glycol butyl ether (EGBE) and methylene chloride can be traced to consumer commodities like paints, pens, and markers (**Figure 7B**). However, certain high and low Z-score chemicals, including some of the leading chemical hits from our screening platform such as methylbenzethonium chloride and tetraconazole lacked available usage category data. Thus, while this resource helped uncover several distinct consumer applications associated with reprotoxic chemicals, primarily cleaning, construction-maintenance, and the consumer goods sectors, the source of their exposure was not comprehensively identified through this database.

**Figure 7:**
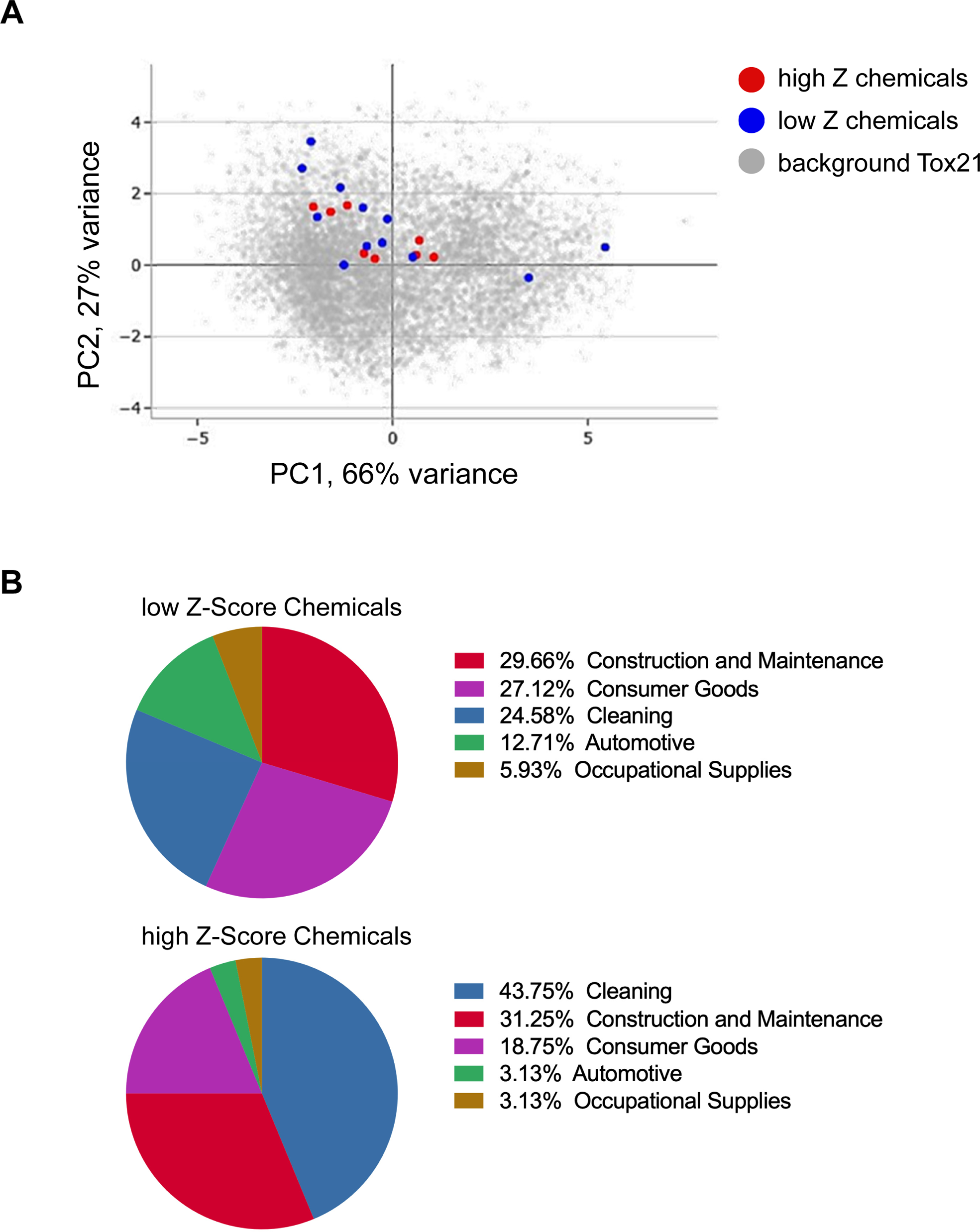
Mining of the Integrated Chemical Environment database. (A) Static PCA plot based on chemical properties for the top and bottom chemicals by Z-score in the *C. elegans* assay found in the ICE database. Background represents the data from >9,000 chemicals tested in Tox21. (B). Use categories of top and bottom chemicals identified in the ICE database.

## DISCUSSION

We have previously shown that *C. elegans* can serve as a rapid and efficient model to identify chemicals with reproductive toxicity activity and dissect their mechanisms of action ^9,10,12,14^. In the present study, we adapted a previously developed and validated assay ^14^ and established a high-throughput platform, enabling us to conduct a comprehensive assessment of 130+ chemicals at multiple doses. We identified 13 chemicals which caused an increase in aneuploidy by at least 1 standard deviation above the mean (Z >1) at any of the concentrations tested, with 5 chemicals displaying a Z-score >1 at 2 or more concentrations (**Table 1**). Two chemicals, benzyldimethyldodecylammonium chloride, a QAC commonly used as a preservative agent and disinfectant ^28^, and methylbenzethonium chloride, a QAC and broad spectrum antimicrobial agent and preservative used in consumer product and also as treatment for leishmaniasis ^29,30^, generated a Z-score >1 at 3 or more of the concentrations tested. Several hits in this screen had already been identified in our initial fully manual version of the screen such as bisphenol A, diazinon, dicofol, norflurazon, thiabendazole, and triflumizole ^14^. Finally, 4 out of the top 5 chemicals further tested for reproductive phenotyping, including the two aforementioned QACs, also displayed a remarkable increase in embryonic lethality, reaching >20% lethality at the test concentration of 100 μM.

There is little mechanistic information regarding the screen’s top hit, methylbenzethonium chloride, that could illuminate why it elicits such remarkable reproductive toxicity in *C. elegans*. As a quaternary ammonium salt, its physicochemical properties, *i.e.* its positive charge, allows it to prevent bacterial attachment to surfaces and therefore has broad-spectrum activity against both gram-positive and gram-negative bacteria. In a study comparing bioactivity signatures obtained from the Tox21 dataset, methylbenzethonium chloride was ranked as one of the top 30 most bioactive chemicals ^31^. Finally, a study examining the effects of structurally-related QACs on reproduction in mice reported a negative impact on key measures of fertility and fecundity such as an increased time to first litter, longer pregnancy intervals, and a reduction in the number of pups per litter and pregnancies ^32^. In a follow-up study, male and female reproductive processes were examined in more detail revealing a significant effect of QACs on various aspects of reproduction (e.g. estrous cycle, gestation, sperm number and motility, etc) ^33^. However, there is reproductive data paucity for the aforementioned top 2 chemicals identified in our screen which are also quaternary ammonium compounds. Thus, our results suggest that QACs may be prioritized for in-depth investigation of their impact on reproductive function across species.

Despite the high bioactivity of methylbenzethonium chloride, there was remarkably no significant difference in overall bioactivity, as measured by the proportion of active to inactive assays (**Figure 4**) and by mean AC50 (**Figure 5**) between the top and bottom chemicals identified in our screen. However, the distribution of assay categories for the ToxCast assays with positive hit calls between the high and low Z score chemicals was significantly different. However, it is notable that the analysis of the intended target families identified “steroid hormones” as a major difference in hits between the two chemical groups. Indeed, the *C. elegans* genome contains a remarkably high number of nuclear hormone receptors and endocrine disruptors have been shown to alter germline function in the nematode in ways that are comparable to mammals ^10,14,34–36^. Further work may examine the genes and pathways underlying the categorical differences which may point towards the mechanisms underlying the reproductive toxicity outcome. Nonetheless, based on the set of chemicals compared in this study, overall bioactivity in the current ToxCast assays may not be a useful metric to prioritize chemicals for reproductive toxicity assessment.

The present study presented here directly compared the outcome *C. elegans* assays (green eggs and him assay, plate phenotyping) with mammalian *in vitro* assays mined from the ToxCast data. Thus, one limitation of the work is the absence of direct comparison between the *C. elegans* results and equivalent mammalian outcomes. One source of this limitation is the relative paucity of mammalian reproductive toxicity data, including from *in vivo* endpoint databases such as ToxRef ^37,38^. This data paucity is apparent in terms of the limited number of chemicals that populate ToxRef compared to other databases but also in terms of the chemical space explored where QACs are underrepresented ^39^. Importantly, reproductive studies also rarely include an in-depth examination of the earlier stages of germline development, such as meiotic differentiation, despite evidence that this period can provide a window of sensitivity to environmental exposures ^40,41^. Thus, the use of a model system such as *C. elegans* where molecular readouts can be functionally tied to germline homeostasis may be useful for prioritization in mammalian models.

## ACKNOWLEDGEMENT

The authors would like to thank Robert Damoiseaux and the Molecular Screening Shared Resource (NCI P30CA016042) for support and guidance during the chemical screening process. We thank Nicole Kleinstreuer for guidance on leveraging the ICE database and we also thank Donatello Telesca for input in the statistical analysis of the HTS data.

---

**Table S1. Z score of all chemicals tested**

